# Evidence for widespread cytoplasmic structuring into mesoscopic condensates

**DOI:** 10.1101/2021.12.17.473234

**Authors:** Felix C. Keber, Thao Nguyen, Clifford P. Brangwynne, Martin Wühr

## Abstract

Eukaryotic cytoplasm organizes itself via both membrane-bound organelles and membrane-less biomolecular condensates (BMCs). Known BMCs exhibit liquid-like properties and are typically visualized on the scale of ~1 μm. They have been studied mostly by microscopy, examining select individual proteins. Here, we investigate the global organization of native cytoplasm with quantitative proteomics, using differential pressure filtration, size exclusion, and dilution experiments. These assays reveal that BMCs form throughout the cytosplasm, predominantly at the mesoscale of ~100 nm. Our data indicate that at least 18% of the proteome is organized via such mesoscale BMCs, suggesting that cells widely employ dynamic liquid-like clustering to organize their cytoplasm, at surprisingly small length scales.

---

Subcellular compartmentalization is an essential feature of eukaryotic life, coordinating the myriad reactions inside a cell. Compartments can colocalize cooperating enzymes to increase their efficiency or spatially separate incompatible reactions or molecular species. Eukaryotic cells achieve compartmentalization both via membrane-bound organelles, e.g., mitochondria or nuclei, and via membrane-less organelles, e.g., the nucleolus. Many membrane-less organelles appear to be assembled through liquid-liquid phase separation (LLPS), which results in concentrated liquid-like droplets known as biomolecular condensates (BMCs) [1,2]. BMCs serve a multitude of functions, e.g., in the regulation of transcription [3], RNA management [4], or signaling [5,6]. Likewise, their malfunction can cause diseases [7,8]. Protein components of BMCs have typically been identified via imaging, co-isolation, or proximity labeling [9–14]. However, these approaches require prior knowledge of at least one constituent of the assembly. Moreover, imaging approaches favor the detection of large (~1 μm) assemblies, due to the diffraction limit of light microscopy, and are often facilitated using overexpression of labeled proteins, with potential impacts on native condensate structure. To date, these approaches have identified about ~100 “scaffold” proteins suggested to drive LLPS [15], which is ~0.5% of the human proteome. However, based on sequence similarity to these proteins, it has been speculated that as much as 20% of the proteome is functionally involved in LLPS [16]. It remains unclear to what extent the cytoplasm is organized into condensate-like structures and at what length-scale these typically form.

To assay the physical properties of protein assemblies throughout the native cytoplasm, we sought to combine filtration experiments of undiluted cytoplasm with quantitative proteomics. We reasoned that we could identify BMCs based on their behavior upon filtration with various pressures. When encountering a pore, assemblies smaller than the pore diameter will pass freely. However, the permeation of assemblies larger than the pore will depend on their deformability, and the applied pressure. Large solids will not pass, regardless of applied pressure, whereas Young-Laplace theory predicts liquid assemblies to pass, if the transmembrane pressure exceeds a critical value [17](Fig. 1A). This is supported by proof-of-principle in-vitro experiments with an imaging assay (fig. S1). To study the organization of intact cytoplasm, we chose lysate from eggs of the frog *Xenopus laevis*, which provides easy access to large amounts of undiluted cytoplasm, in a near-native state. We verified that the extract preparation spin used to crush the eggs did not sediment proteins known to be involved in BMCs (fig. S2). Cytoplasmic extract was centrifuged against polyethersulfone filter membranes with a defined particle size cutoff diameter (*d_pore_*). We analyzed the filtrates of various experimental conditions by multiplexed proteomics, quantifying each protein’s concentration relative to the input (Fig. 1B) [18–20].

**Figure 1:**
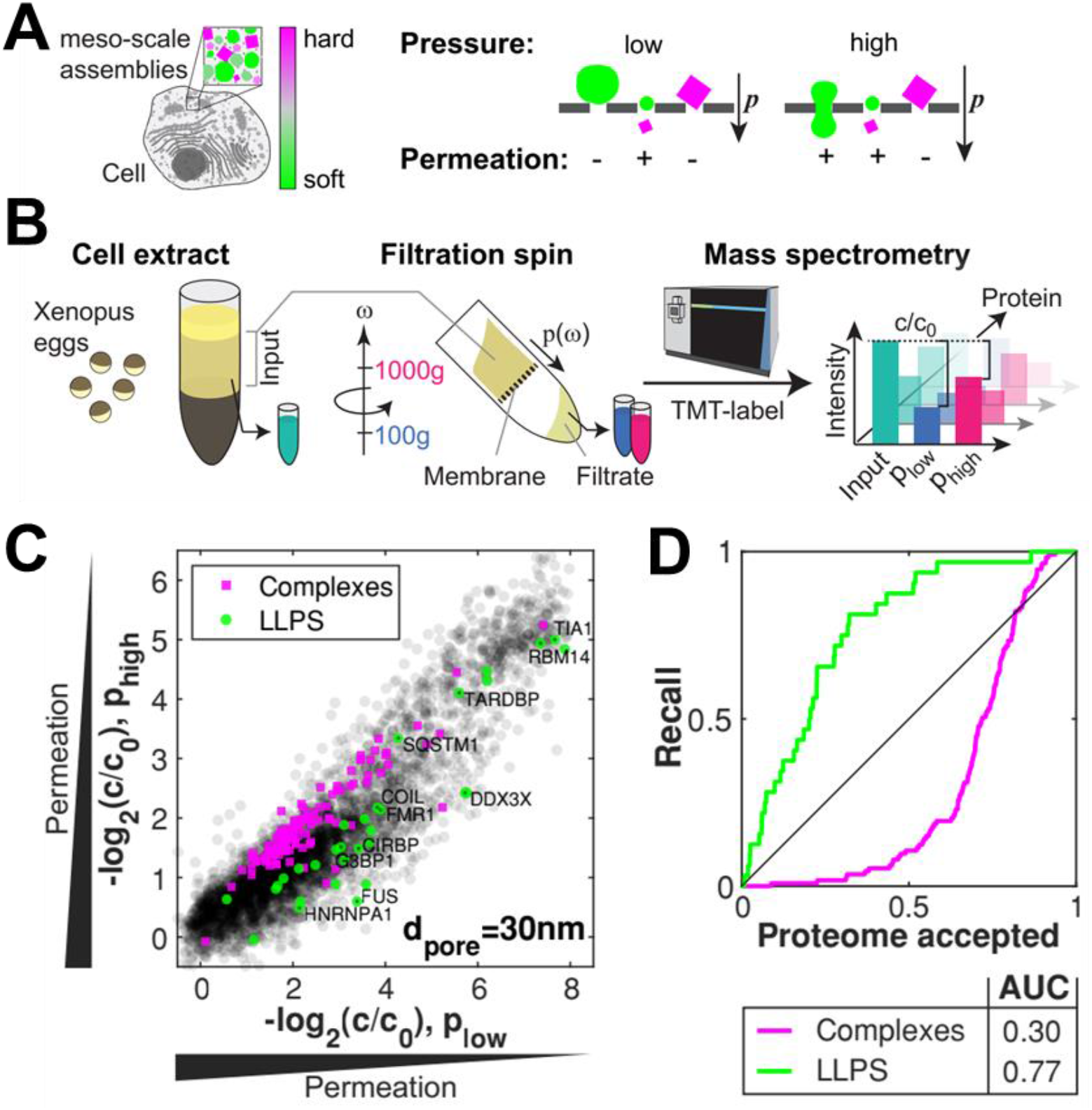
Proteomics of differential pressure filtrated cytoplasm reveals organization in liquid-like assemblies. **A)** Illustration of the pressure dependence of soft (green) and hard (magenta) assemblies in filtration experiments. **B)** Experimental outline. We spun undiluted cytoplasm from frog eggs through filters at different pressures (*p_low_* at 100g and *p_high_* at 1000g). We analyzed the filtrate in comparison to input by mass spectrometry. For each protein, we obtain the filtrate concentration relative to the input concentration via quantitative proteomics c/*c*_0_. **C)** Scatter plot of protein permeation at high versus low pressure. Close to the diagonal, the permeation is pressure independent; below the diagonal permeation is pressure sensitive. Proteins established to form LLPS (green), or large complexes (magenta) exhibit distinguishable behaviors. **D)** Results of experiments at different pore sizes (30 nm, 100 nm) are integrated into one metric for squeezing behavior and used to order the proteome. The resulting receiver operating characteristic (ROC) preferentially recalls LLPS proteins while complexes are underrepresented.

We compared the permeation of all proteins at fixed pore size (*d_pore_* = 30 *nm*), but with varied pressure, applied by varying the centrifugation speed (*p_low_* at 100g and *p_high_* at 1000g). We observe a broad spectrum ranging from free passage to heavy retention (Fig. 1C), which can be categorized into two regimes: 1) A large part of the proteome shows approximately pressure-independent permeation (close to diagonal) as would be the case for both large solid-like complexes and assemblies that are smaller than the pores (close to origin). 2) For many other proteins, permeation is increased at higher pressure (below the diagonal), as would be expected for deformable, liquid-like assemblies. Consistent with these predictions, annotated canonical large protein complexes (magenta) reveal a preponderance along the diagonal, while proteins known to be associated with LLPS (green) [21–25]) (table S1) are enriched below the diagonal, i.e., they exhibit enhanced elution at higher pressures. Executing the experiment with a larger pore size (*d_pore_* = 100 *nm*) yields qualitatively similar results (fig. S3). To characterize overall squeezing behavior, we merge the results for different pore sizes (*d_pore_* = 30 *nm*, 100 *nm*) by summing each protein,s distance to the diagonal. This “squeezing score” displays significant differences between known LLPS proteins and the whole proteome (Fig. 1D). These findings suggest that differential pressure probes mesoscopic physical properties of the cytoplasm and could be used to identify novel BMCs and their components.

A characteristic property of phase-separated BMCs is that they typically form via dynamic, multivalent interactions and disassemble below a particular saturation concentration [1,2,26,27]). To test whether altering concentration impacts the apparent mesoscopic cytoplasmic organization, we diluted cell extracts to various extents and examined their filtration behavior at fixed pore size (*d_pore_* = 100 *nm*) and pressure *p_low_* (fig. S4A). Remarkably, at a dilution of only 1.4-fold, known LLPS proteins show significantly different permeation behavior compared with canonical larger complexes or the entire proteome (fig. S4B). LLPS proteins exhibit a progressive increase in permeation with dilution, that likely originates from shrinking condensate sizes as concentrations approach and cross below different saturation thresholds.

We next sought to integrate all our measurements to improve predictions on which proteins are associated with phase-separated condensates. Existing predictors of proteins driving LLPS analyze published data to extract sequence features but suffer from a lack of comprehensive reference data. To this end, we trained a classifier to identify LLPS proteins (Fig. 2A). The classifier learns by bagging an ensemble of linear discriminators and decision trees [28–30]. As features, we used the results from our differential pressure filtration and dilution experiments. We additionally included coarsened sequence information on intrinsic disorder [31], nucleic acid-binding [32–34], and amino acid composition [33]. The performance of our predictor is assessed by the recall of known LLPS proteins with 5-fold cross-validation. While prediction using experiments alone already has an area under curve (AUC) of 0.86, our final predictor, including all features, reaches an AUC of 0.93 (table S2). This approach establishes an improvement over the state-of-the-art predictions of LLPS proteins by catGRANULE (AUC=0.84, [35]), Pscore (AUC=0.85, [36]), or PSAP (AUC=0.88, [22]). Notably, we gain sensitivity by a steeper early increase, which is arguably the most relevant regime for the prediction (Fig. 2B).

**Figure 2:**
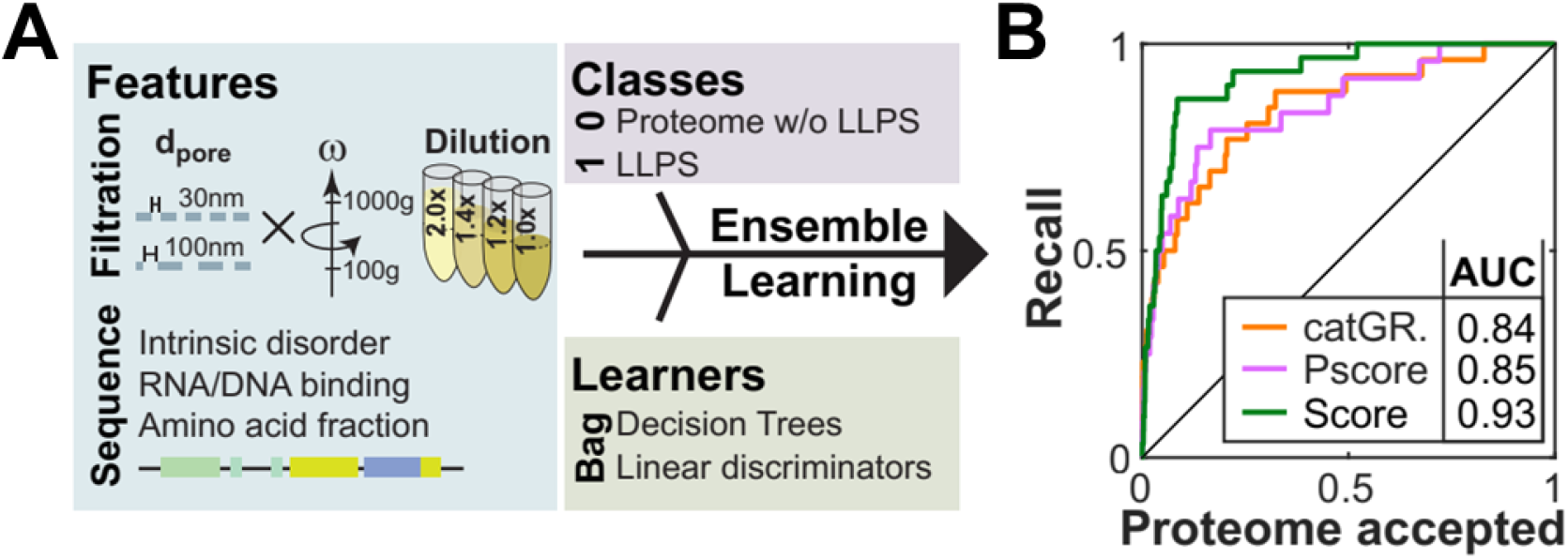
Integrating proteomics experiments enhances LLPS prediction. **A)** Classification ensemble learning to distinguish LLPS proteins from the bulk proteome. We use the experimental data and additional sequence annotations as features. Bagging of decision trees and linear discriminators returns the classification score. **B)** ROC displaying the recall of LLPS proteins by the respective scoring. The AUC of our prediction score (dark green) is higher than the one of the established predictors catGRANULE (orange, [35]) and Pscore (purple, [36]) and shows a steeper increase.

Cellular BMCs to date have been reported mainly on the micrometer length scale. To investigate the length scale of the liquid-like organization in our system, we compared the elution behavior at different pore diameters (30/100/200 nm). At the largest pore size of 200 nm, we observe an overall higher permeation with very few proteins retained (Fig. 3A). Notable exceptions include the unsurprising retention of mitochondrial proteins (Fig. 3A). The striking transition in the behavior of cytoplasm as the pore size is decreased to 100 nm, suggests this is a characteristic length scale of liquid-like organization. This is further supported by filtration experiments in a complementary setup, using gravity flow on polymer mesh filters (fig. S5). These data suggest the widespread presence of liquid-like cytoplasmic assemblies on the sub-micrometer scale, involving a broad swath of the proteome.

**Figure 3:**
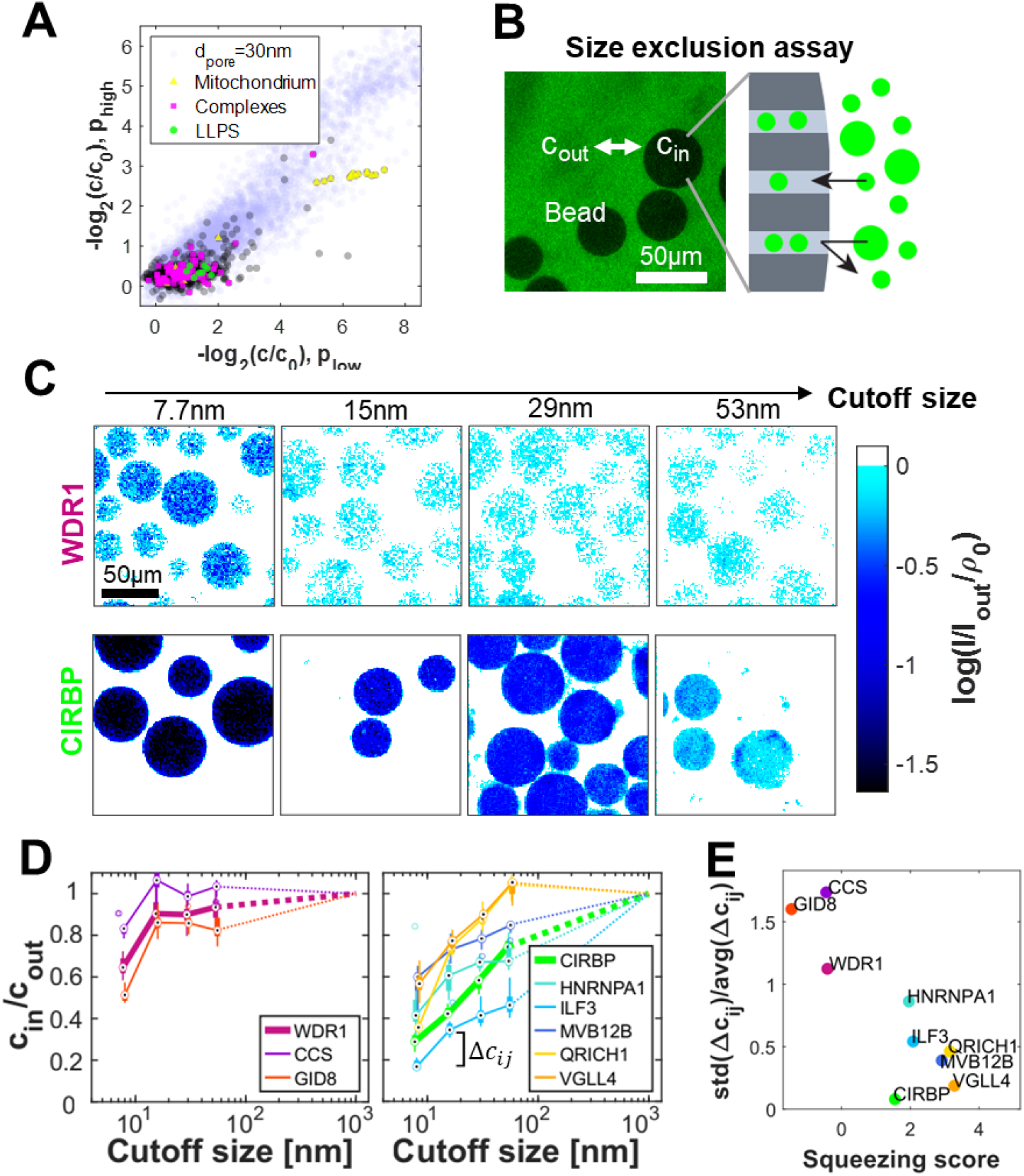
Proteomics and imaging assays indicate that liquid-like assemblies predominantly organize at the ~100 nm length scale. **A)** The filtration experiment with 200 nm pore size indicates that few proteins besides mitochondrial ones (yellow) are retained (30 nm data for comparison (lilac)). **B)** Microscopy of GFP-labeled proteins with size exclusion beads to assess protein assembly sizes. The concentration inside a bead’s polymer matrix *c_in_* compared to outside solution *c_out_* provides information about the protein’s assembly size. The example image shows beads with 400kDa size cutoff in an extract expressing CCS-GFP. **C)** Examples of size exclusion assay for the proteins WDR1-GFP and CIRBP-GFP (examples of proteins in small assemblies and LLPS respectively) with four different bead cutoff sizes. Confocal image intensities *I* are normalized to their respective outside solution intensity *I_out_* and each column is scaled by the accessible volume for monomers *ρ*_0_ of the used bead. Thus, white bead color stands for *c_in_/c_out_* ≈ 1 and darker blue tones report higher exclusions. **D)** Size-exclusion plots. Proteins devoid of squeezing behavior (left panel) show a characteristic jump at short length scale. In contrast, proteins filtrations suggested to be in BMCs (ILF3, MVB12B, QRICH1, VGLL4) including well-established LLPS (CIRBP, HNRNPA1) show organization throughout the meso-scale. Dotted lines extrapolate data to 1 at 1μm based on lack of visible organization via microscopy. **E)** Scatter plot of observed assembly behavior versus squeezing score (Fig. 1D). We observe organization over wide length scales for proteins that are predicted to be in liquid-like assemblies.

We next sought to validate our findings of liquid-like organization on surprisingly short length-scales using an experimentally orthogonal light microscopy assay, that investigates the size of cytoplasmic assemblies below the diffraction limit. We selected proteins exhibiting different filtration behavior and expressed GFP-tagged versions by doping the cytoplasm with the corresponding mRNA. Interestingly, at our low expression levels of only a few nM, we can detect fluorescence, but none of the investigated proteins - including established LLPS proteins like HNRNPA1 [37] or CIRBP [38] - show any signs of assemblies on the micron-scale (fig. S6). To interrogate potential assemblies on smaller length scales, we equilibrated the lysates with size exclusion beads (Fig. 3B). The size exclusion of biomolecular assemblies by a bead’s polymer matrix reduces the concentration inside the bead *c_in_* compared to the outside *c_out_*. The concentration ratio *c_in_/c_out_* at different cutoff sizes (~7.7/15/29/53 nm) serves as a proxy for the cumulative assembly size distribution. We measure *c_in_* and *c_out_* by their GFP intensity and correct *c_in_* for the excluded volume in the beads, which we determined with a dextran-rhodamine solution (hydrodynamic radius of ~5nm) (Fig. 3C and fig. S7). Proteins that showed pressure-independent permeation in the filtration (WDR1, GID8, CCS) mainly organized on the 10 nm scale. By contrast, proteins that we found in the pressure-dependent regime (MVB12B, VGLL4, QRICH1, ILF3) exhibit increasing *c_in_/c_out_* with the cutoff size, similar to established LLPS proteins (CIRBP, HNRNPA1) (Fig. 3D, E). This behavior suggests that the size of these assemblies is not sharply defined but spans across the sampled scale, matching the expectation for phase-separated droplets, which appear to assemble typically on the scale of ~100 nm.

We next wanted to estimate what fraction of the proteome organizes into such mesoscale liquids. To this end, we developed a noise model based on replicates for our integrated differential pressure filtration and dilution experiments (Fig. 4A). 22% of the proteome falls beyond a 2% false discovery rate, including 68% of the LLPS references (Fig. 4A). Additionally, only 12% of the detected proteins do not respond to the filtration and appear unstructured at the assayed scale. Database annotations assign ~40% of the proteins to membrane-bound organelles other than the nucleus. Together, we conclude that at least 18% of the detected proteins are in BMCs (Fig. 4B). However, based on the spread of known BMCs into our noise model we believe this is a very conservative lower bound (Fig. 4A). We expect a significant fraction of the remaining third of the proteins to be also organized via BMCs (Fig. 4B).

**Figure 4:**
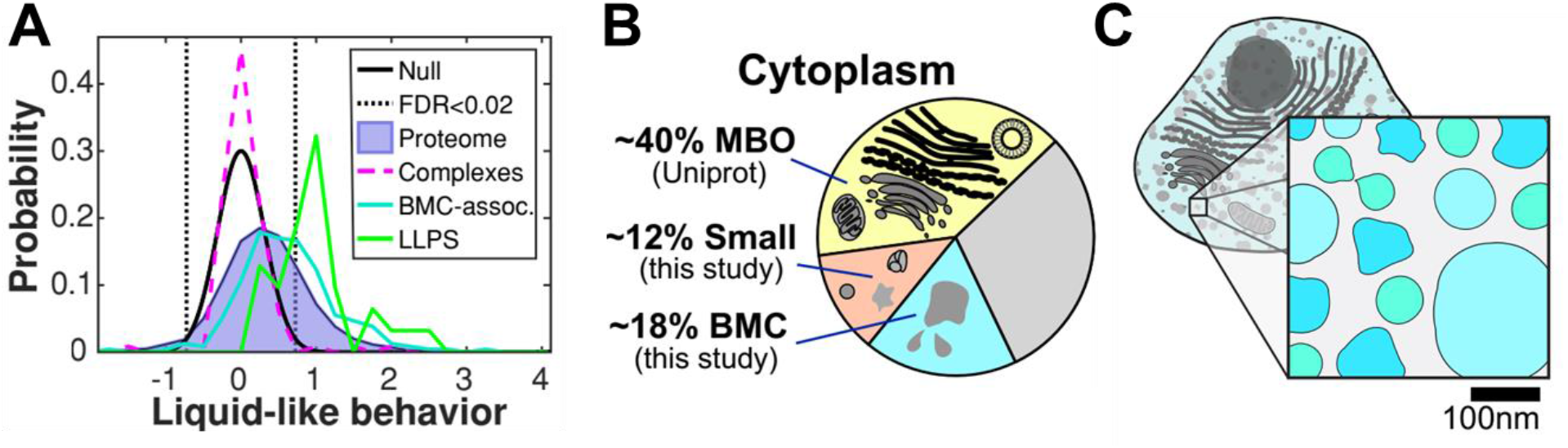
At least 18% of the cytoplasm organizes in BMCs. **A)** We define liquid-like behavior as the accumulative shift in dilution and differential pressure filtration experiments, where the null-model (black) is the replicate noise around strongly retained assemblies (complexes (magenta) and transmembrane domain proteins). Beyond a 2% false discovery rate (black dotted lines), we find 22% of the proteome (blue, filled) and 68% of the LLPS references (green). BMC-associated, supposably not LLPS driving, proteins (teal) are also shifted towards this regime. **B)** Cytoplasmic organization to a large extent is achieved by membrane-bound organelles (~40%, yellow, Uniprot). In our filtration experiments, only around 12% of proteins are not affected (salmon). Based on the 2% FDR, 18% of the cytoplasm is organized in BMCs (cyan). We currently cannot confidently assign organization modality for the remaining third of the proteome (gray). **C)** Our results suggest that biomolecular condensates contribute significantly cytoplasmic organization at the ~100nm scale. This indicates that cytoplasm is widely structured at the mesoscale.

Previous studies have focused primarily on large (≥1μm) condensates, which are easily accessible by microscopy. Our differential filtration and size exclusion studies on intact cytoplasm reveal that phase-separation prone proteins ubiquitously organize cytoplasm into smaller, “mesoscopic” assemblies, which exhibit pressure-dependent, liquid-like deformability (Fig. 4C). It remains an exciting question how these tiny BMCs can exist without coalescence or Ostwald ripening, i.e., the expected growth of large condensates at the expense of smaller ones. We speculate that this mesoscopic organization is highly dynamic, reflecting continuous assembly and disassembly. Such behavior can originate from chemical activity [39], but is also reminiscent of phase separating systems in the vicinity of a critical point, as has been suggested for two-dimensional phase separation in the plasma membrane [40].

## Acknowledgments

We thank Lillia Ryazanova for help with sample preparation and members of the Wühr and the Brangwynne Laboratories for useful discussions. We thank David Hill for access to the Xenopus ORFeome and James Pelletier for help with designing the filter holders. This work was supported by EMBO ALTF 601-2018 (FCK), the Lewis-Sigler Institute (MW, CPB) NIH grant R35GM128813 (MW), American Heart Association predoctoral fellowship 20PRE35220061 (TN), Princeton Catalysis Initiative (MW), Eric and Wendy Schmidt Transformative Technology Fund (MW, CPB) and the Howard Hughes Medical Institute (CPB).

## Author contribution

FCK, CPB, and MW designed the research. FCK conducted the experiments and analyzed the data. FCK and TN developed and performed in vitro protein expression. MW and CPB provided funding and supervised the study. FCK, CPB, and MW wrote the manuscript, and all authors helped edit the manuscript.

## Supplementary Figures

**Supplementary Figure S1:**
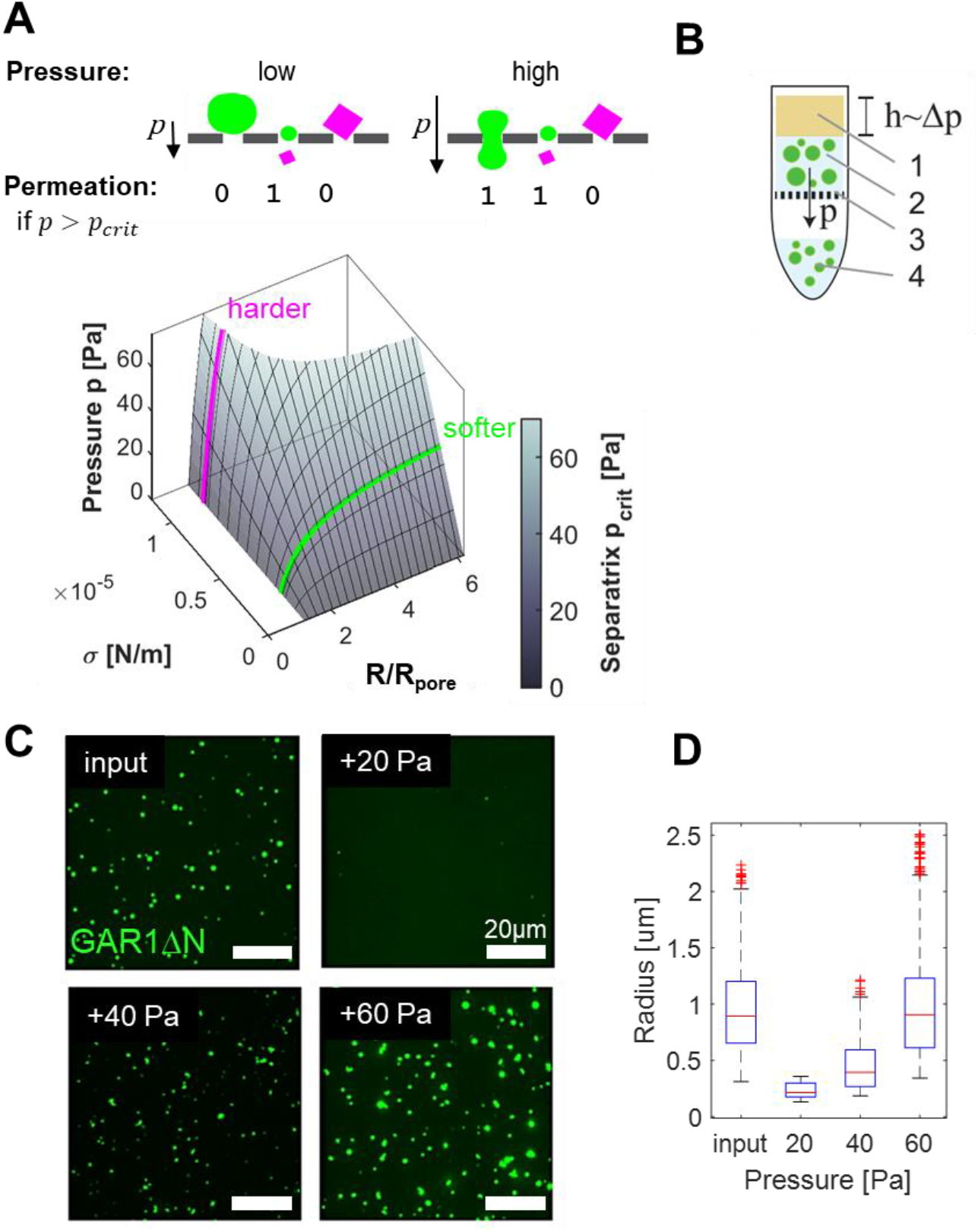
Differential pressure filtration allows differentiation of liquid-like or solid assemblies. **A)** Schematic illustrating the pressure dependence of softer (green) and harder (magenta) assemblies. Young-Laplace based theory predicts distinct regimes of retention and permeation of liquid droplets [17]. Theory parameters are the droplet radius R relative to the pore radius Rpore, the surface tension σ, the contact angle (not shown), and the transmembrane pressure *ρ* In the R-σ-p space, these regimes are separated by the surface of critical pressure *p_crit_*. Softer assemblies squeeze through pores bigger than themselves at higher pressures, while harder assemblies are almost only sensitive to size. **B)** In-tube filtration setup. An in vitro solution of phase-separated GARI△N-GFP droplets flows through a porous membrane (polyethersulfone, nominal size cutoff 0.8 μm). Gravity drives the flow, and the applied pressure is varied by the height of a layer of mineral oil. **C)** Fluorescence micrographs corresponding to the experiment described in B, increasing the transmembrane pressure by 20, 40, and 60 Pa. Bars are 10um. **D)** Pressure-dependent droplet size distribution. Resolution is achieved at low pressures, while higher pressure results in total permeation of the input.

**Supplementary Figure S2:**
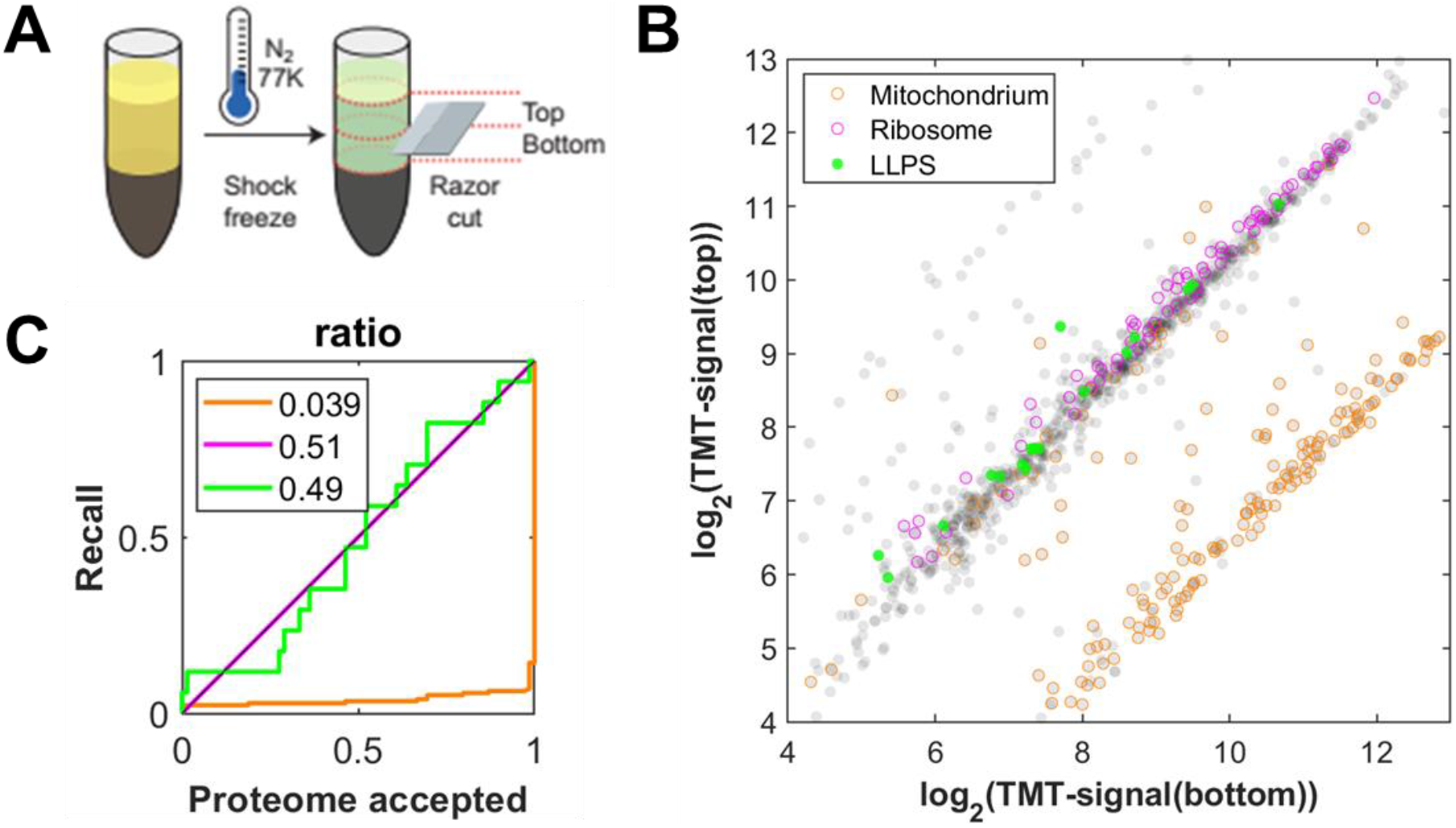
Sedimentation in extract preparation spin is neglectable. **A)** Schematic illustrating the sample collection. After 16min of the preparation spin (14400g), the tube is shock frozen in liquid nitrogen and subsequently cut into a top and a bottom section with a razor blade. **B)** Comparison of the two halves of the extract. While mitochondrial proteins (orange) are shifted to the bottom, indicating sedimentation, the bulk part of the proteome, including both ribosomal (magenta) and LLPS proteins (green), stays unchanged. Scatter plot of raw TMT signals. Text labels for LLPS proteins. **C)** Receiver operator characteristics on the signal ratio top to bottom of the data in B.

**Supplementary Figure S3:**
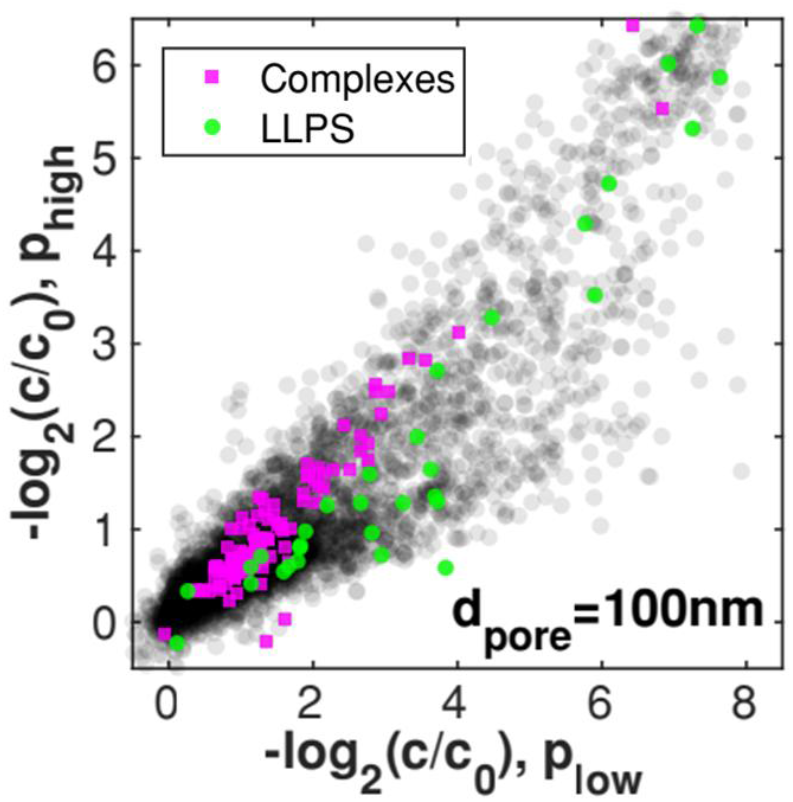
Differential pressure filtration at pore size 100 nm. Executing the experiment presented in Fig. 1C with a larger pore size (*d_pore_* = 100 *nm*) yields qualitatively similar results. However, there are qualitative changes and, as expected, the overall permeation is higher.

**Supplementary Figure S4:**
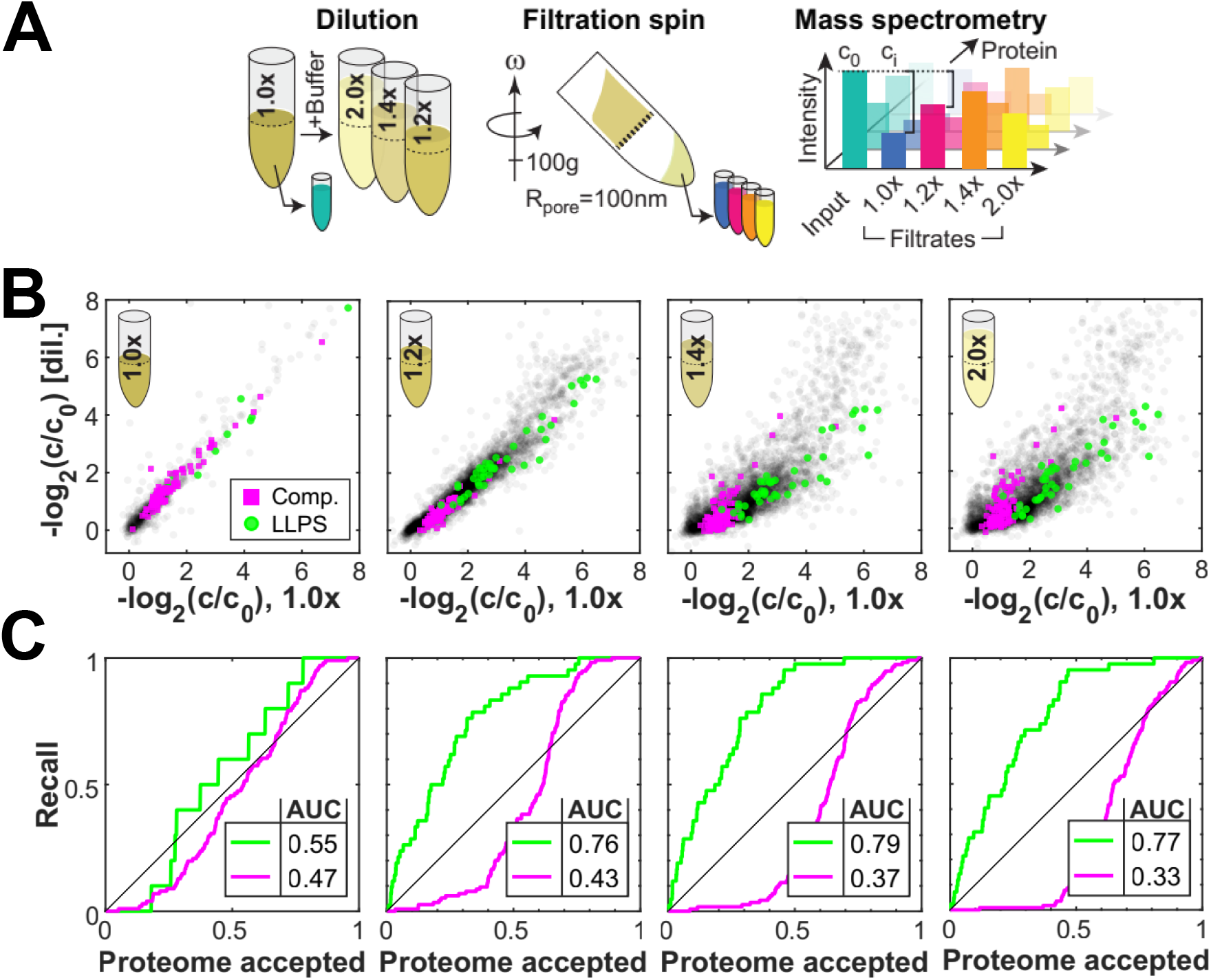
Liquid-like assemblies are sensitive to moderate dilution. **A)** We performed the cytoplasmic filtration experiment with 100 nm pores and low pressure (100 g) and compared protein retention with lysate diluted by various amounts. The concentrations of the filtrates relative to the input are measured by mass spectrometry. **B)** Experiment of filtration of diluted lysate reveals that LLPS proteins seem to disassemble at dilutions larger than 1.2x. **C)** Receiver operating characteristics to the plots above in (B) for the ratio of the permeated concentrations *c_y_*/*c_x_*, where *c_x_* and *c_y_* are the concentrations of undiluted and diluted condition, respectively.

**Supplementary Figure S5:**
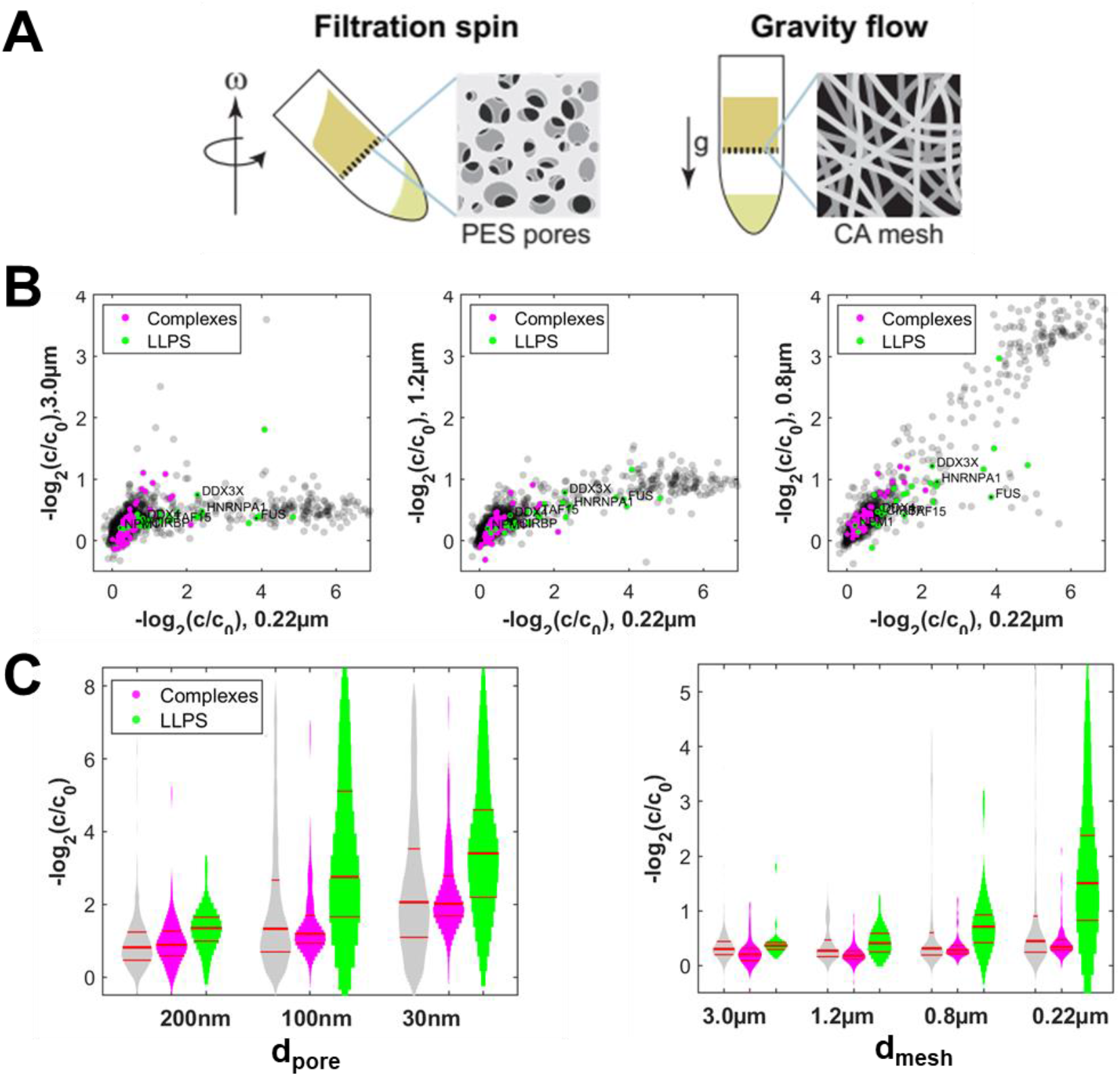
Pore size dependence of filtration. **A)** Schematics of the spin filtration setup, using polyethersulfone (PES) membranes and the alternative setup gravity flow setup, using cellulose acetate (CA)-mesh filters. The large open area of the CA mesh enables flow through at 1g force. **B)** Scatter plots of the CA gravity flow experiments. Larger meshes (3 um, 1.2 um, 0.8 um; left, mid, right panel on y-axis) can only resolve few structures compared to a 0.22 um mesh (x-axis). LLPS proteins are shifted to less permeation. **C)** Permeation histograms of the PES filters at low pressure (from main text) (left) and the CA mesh filters (right). The retention increases with smaller pores or meshes, suggesting assemblies on the 100nm length scale. This behavior is pronounced for LLPS proteins.

**Supplementary Figure S6:**
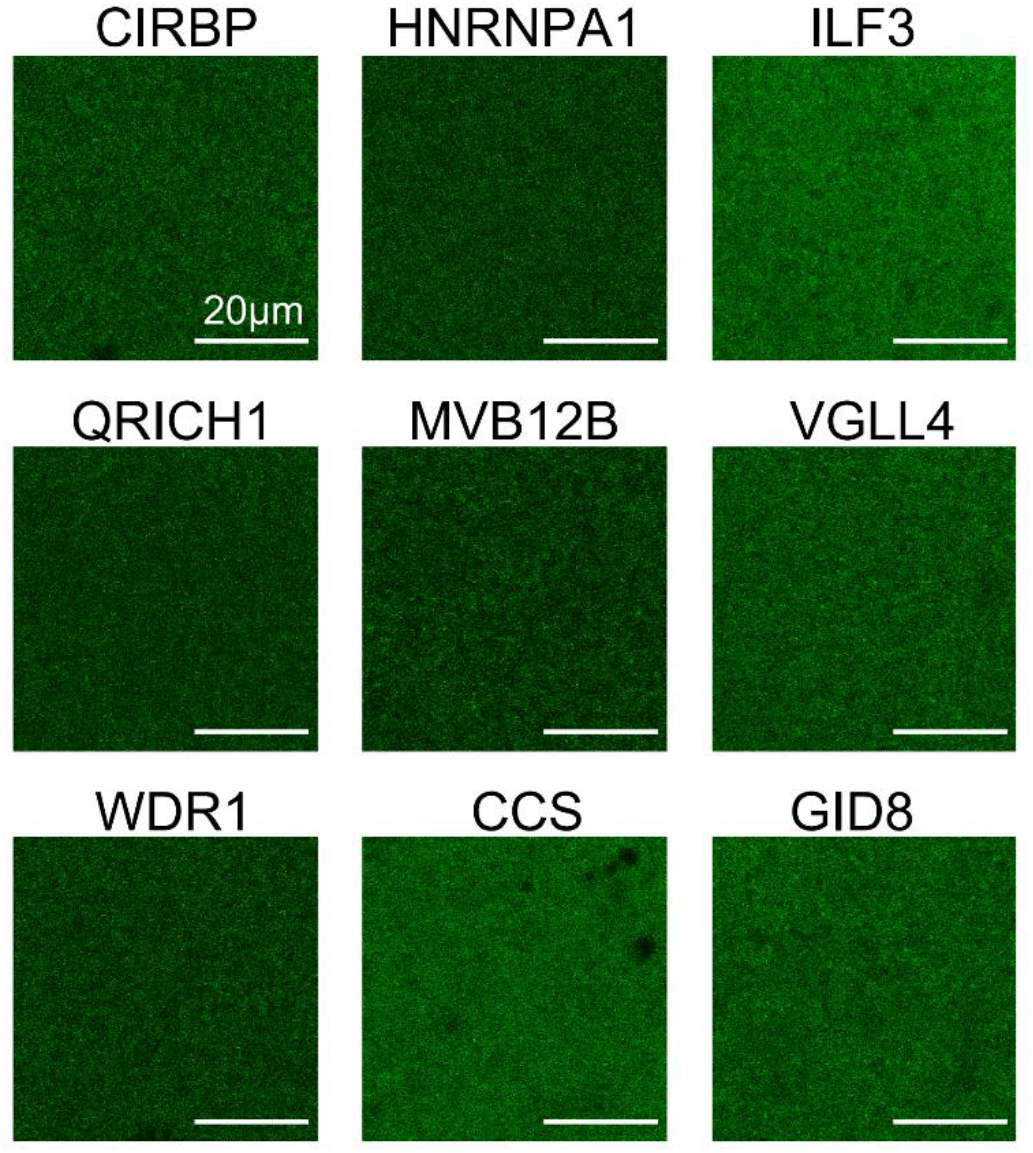
Confocal micrographs of GFP-fused proteins expressed from mRNA in the cell extract. Solutions appear relatively homogenous, and we did not detect structures on the micrometer scale. Lookup tables adjusted individually to 0.35% saturated pixels to enhance contrast.

**Supplementary Figure S7:**
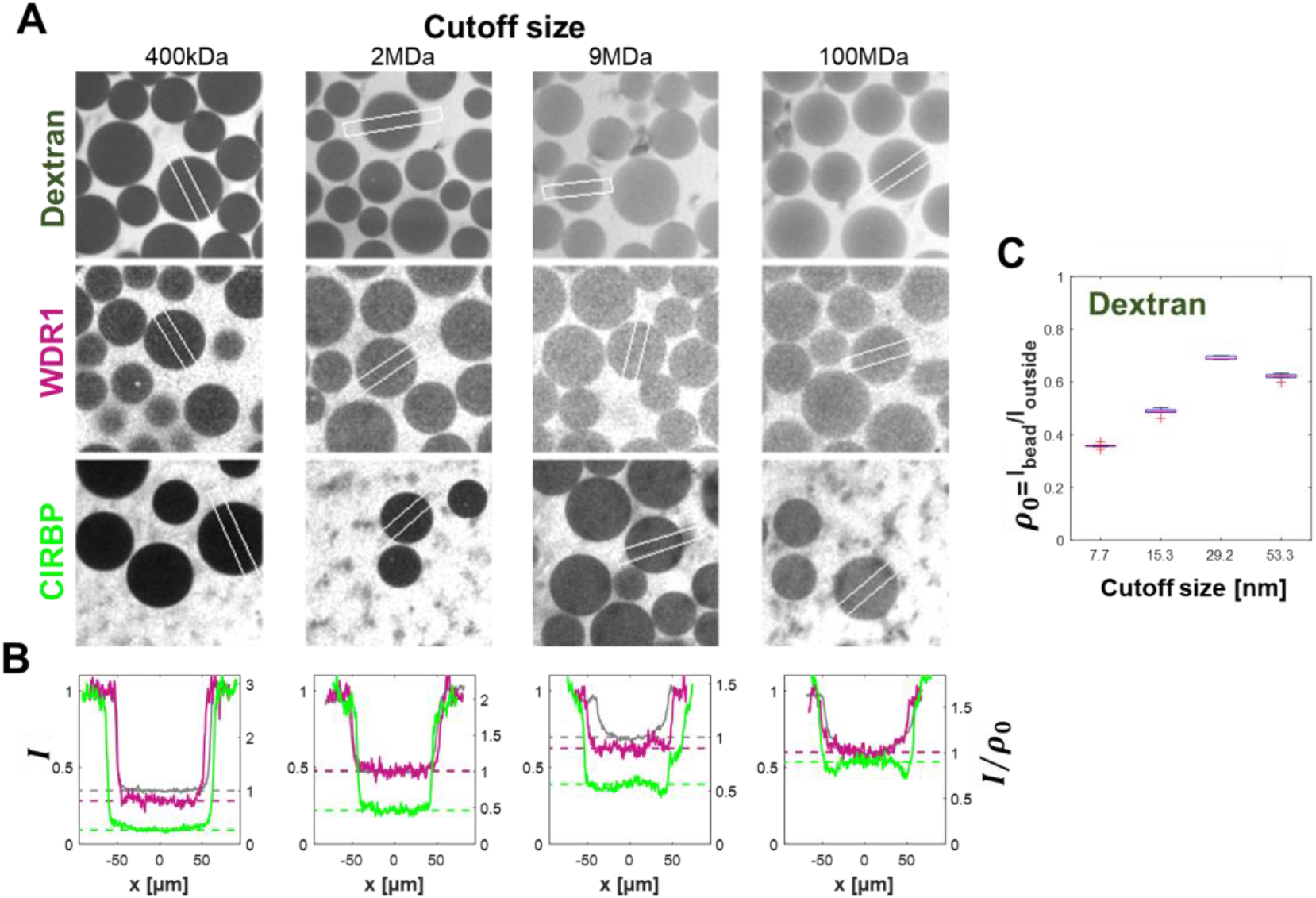
Size exclusion assay. **A)** Confocal micrographs of the size exclusion assay described in the main text. Chromatography beads of different cutoff sizes are placed in extracts with GFP-labeled proteins (WDR1 and CIRBP) and Dextran 70 kDa solution for reference. Top panels display the calibration measurement with dextran 70k Da-rhodamine. Intensities are normalized to the outside solution. White rectangles indicate regions of the line plots in panel B. Intensities are normalized to the outside solutions for comparability. **B)** Density comparison of the solutions above. Left y-axis displays the normalized, background corrected fluorescence intensity *I*. Right y-axis displays the intensity normalized to the dextran density *I*/*ρ*_0_., as done to correct for accessible volume. **C)** Measurement of dextran intensity ratios that serve as normalization of the assay. The bright, homogenous solutions allow for a precise determination of ρ_0_. Boxplots summarize ~7 bead measurements.

