## Supplementary Text for "Evidence for widespread cytoplasmic structuring into mesoscopic condensates"

### Materials and Methods

#### Xenopus laevis egg extracts.

Frog husbandry: Mature *Xenopus laevis* females were purchased from Nasco and maintained by Laboratory Animal Resources at Princeton University. All animal procedures are approved under IACUC protocol 2070, reviewed in April 2021. Ovulation was induced with at least six-month rest intervals.

Egg collection: *X. laevis* eggs were collected as previously described [1]. Female frogs were primed with pregnant mare serum gonadotropin (HOR-272, ProSpec-Tany TechnoGene Ltd) within two months before the experiment. At 16 h before egg collection, frogs were injected with 500 U of human chorionic gonadotropin (Sigma CG10) and kept at 16°C in Marc's Modified Ringer's [2] (50 mM HEPES pH 7.8, 100 mM NaCl, 2 mM KCl, 1 mM $MgCl_2$, and 2 mM $CaCl_2$). We collected eggs the next day in MMR buffer and sorted out pre-activated ones for further use.

Extract preparation: *X. laevis* egg extracts were prepared as previously described [3,4]. The eggs were dejellied in MMR with added L-cysteine (2 wt%, pH 7.8) and washed in CSF-XB buffer (100 mM KCl, 20 mM HEPES, 2 mM $MgCl_2$, 0.1 mM $CaCl_2$, 4 mM EGTA, pH 7.7). Eggs were collected into 3.5 ml centrifuge tubes (Beckmann) in the presence of cytochalasin D (Sigma C8273) and LPC protease inhibitor premix (leupeptin (Sigma L2884), pepstatin (Sigma P5318), chymostatin (Sigma C7268)). The surplus buffer is removed after a soft spin at 500 g for 1 min. Eggs are crushed and fractionated in a spin at 14400 g for 15 min. The cytoplasmic fraction is extracted using an 18G gauge needle. The extract was supplemented with 10 µg/mL cytochalasin D, 10 µg/mL LPC, 1 µM nocodazole (Sigma M1404), and 50 mM sucrose. All used drugs were dissolved in DMSO, resulting in a total DMSO concentration of <2.5 permille. We prefiltered extracts through a 6 µm polyether mesh filter to remove residual debris and stored them on ice for further use.

#### Preparation spin control experiment

Immediately after the preparation spin, the centrifuge tube containing the spin-crushed and sedimented eggs is shock-frozen in liquid nitrogen at 77 K. The extract section, excluding lipid and yolk/debris sections, is cut out and cut in halves using a razor blade.

#### Filtration experiments

Filtrations are performed in 2 ml tubes in a tabletop centrifuge, using 3D printed filter holders[5]. The design files are available on our GitHub page:
https://github.com/wuhrlab/3DFilterHolderDesigns.

We used the Hubs platform (https://www.hubs.com/) to print the holders using either Standard Resin (SLA) or Dental resin (SLA) materials at 20% infill rate, 50 µm layer height. We use polyethersulfone membranes (Sterlitech, PES00347100, PES0147100, PES0247100, PES0847100) in the spin filtration setup and the in vitro assay, and cellulose acetate membranes (Sterlitech, C080A047A, C300A047A, CA1247100, CA0247100) in the gravity flow setup. Membranes were wetted with methanol and washed with CSF-XB buffer. After excess buffer removal, the sample chamber was flushed twice with 50 µl cell extract.

Differential pressure filtration experiments: 50 µL of the cytoplasmic extract was loaded and spun at the respective speed ($p_{low}$ at 100 g, $p_{high}$ at 1000 g) in turns of 4 min until approximately 15 µL of filtrate was collected. After each turn, we refilled the sample chamber input by the estimated flowthrough volume while also stirring by pipetting to avoid cake formation.

Dilution experiments: 30 min before the spin filtration process, the extracts were diluted in CSF-XB buffer in a dilution series to a factor of 1.2, 1.44, and 2.

Gravity flow experiments: 200 µL samples were loaded and placed in a box providing a humid atmosphere until sufficient flowthrough accumulated. The input chamber was stirred occasionally.

In-vitro assay: Droplets were pre-formed in one batch (2 µM concentration of GAR1ΔN-GFP in 150 mM KCl at pH 7.4 (MOPS)) and distributed among the samples (25 µL each) after 10 min. Various amounts of mineral oil were added to adjust the applied pressure. GAR1ΔN-GFP was prepared as previously described [6].

#### Expression of GFP-fusion proteins

The Gateway entry plasmids of desired proteins were retrieved from *Xenopus laevis* ORFeome [7]. The destination vector carrying an EGFP sequence-TEV site-S-tag was purchased from Addgene (pCSF107mT-GATEWAY-3'-LAP tag, plasmid #67618). For the Gateway LR cloning reaction, the entry plasmid, the destination plasmid, and the Gateway LR clonase II enzyme mix (Invitrogen 11791) were combined at the ratios recommended in the manufacture protocol. After the reaction, the expression cloned vector was purified, then linearized using restriction enzymes, which were chosen so that the region of protein of interest was protected. The linearized plasmids were in-vitro transcribed using the mMESSAGE mMACHINE SP6 kit (Invitrogen AM1340) supplemented with a 7-methyl guanosine cap protected on the 5' end terminal, and a poly(A) tail (NEB M0276). Finally, RNA products were purified using Trizol LS reagent (Invitrogen 10296010), then resuspended in nuclease-free water at ~1 µg/µL in the final RNA concentration. The RNA solution was added in a volume ratio of 1:100 to the extracts.

#### Size exclusion assay

Size exclusion chromatography beads (GE, Sephacryl High Resolution, S-200/300/400/500, 17-584-10/17-599-99/17060999/170613-10; molecular size cutoffs 400 kDa/2 MDa/9 MDa/100 MDa) were washed in CSF-XB buffer and equilibrated in three rounds of sedimentation, supernatant removal, 1:5-add-up in plain cell extract and waiting times of about 10 min. After being finally added to the labeled cell extracts, we waited at least 15 min before imaging. The samples were enclosed in mineral oil to prevent evaporation (Sigma M5904). For calibration, we added the beads to a 70kDa-Dextran-rhodamine-B-isothiocyanate (Sigma, R9379, approx. 5 nm) solution in CSF-XB buffer. Sizes were estimated from molecular weights using Zetasizer (Malvern Panalytical).

Image analysis: Line profiles of beads and surrounding solution were measured manually in ImageJ/Fiji [8,9]. Raw image intensities were corrected for the detector background to make them proportional to concentrations. An estimation of the accessible volume for each bead type was measured by the intensity ratio for the dextran solution. The GFP intensities measured inside the bead were adjusted by this factor to calculate the concentration ratio.

#### Microscopy

Confocal images of labeled cytoplasmic extracts were taken in glass-bottom well-plates (Cellvis) on a Nikon A1 laser scanning confocal microscope using 60x and 20x oil immersion objectives. Images of the in vitro assay were taken on a Nikon spinning disc confocal microscope with a 100x oil immersion objective.

#### Reference databases

If not stated otherwise, all annotations are from Uniprot [10]. Our reference group for large 'complexes' includes proteins from ribosome, proteasome, and the vault complex. In the group 'mitochondrium', we exclude proteins with promiscuous subcellular location annotations. The reference group 'BMC-associated proteins' is the union of the categories' client' and 'regulator' in DrLLPS [11]. For the estimation of the fraction of proteins in membrane-bound organelles, we use QuickGO [12] and restrict to ‘UniprotKB’(swiss-prot) and‘located in‘ membrane-bound organelles. LLPS database: To create a comprehensive, high-confidence database of known LLPS proteins with minimal personal curation bias, we follow the approach of merging several published databases [13]: PhasePro [14], PhaSepDB [15], DrLLPS [11], LLPSDB [16], and extend this set by the reference list for the PSAP predictor [17]. We furthermore include proteins assigned 'candidate' in PhasePro and the updated annotation 'PS-self' in PhaSepDB version 2 that indicates de-novo phase separation without partners. We define the 'consensus level' as the number of databases in which a protein is found; throughout the manuscript we use consensus level 4, except fig. S4 (level 3). Overall, proteins with higher consensus level showed more significant shifts. This may reflect the increasing curation quality but may also be due to system specific or partner dependent phase separation.

#### Predictor learning

We train our predictor for phase separating proteins in Matlab (MathWorks), using the function fitcensemble for ensemble classification. Class one is proteins from our LLPS database and class two is the rest of the proteome. We use only proteins identified in both the differential pressure filtration and the dilution experiments (N~4000) and thus have no missing values. The features are the filtrate concentrations relative to the input in the different conditions (differential pressure filtration: 30 nm/100 nm at $p_{low}$ / $p_{high}$. Diluted filtration: 100 nm at $p_{low}$, diluted 1x (undiluted)/1.2x/1.4x/2x). We include features for intrinsically disordered regions determined by Espritz [18](fraction of amino acids (aa) in IDRs, number of IDRs of >50 aa/>30 aa/any-length, setting: disprot, BestSw). We include features on DNA binding [10], RNA binding (QuickGo [12]) and RNA-binding domains [19]. Following van Mierlo et al. [17], we also include the sequence fractions of Glycine, Cysteine, Leucine, Isoleucine, as well as the content of aliphatic and aromatic residues. We use the method 'Bag' to train ensembles with 500 learners of decision trees and linear discriminators [20-22]. The trees are restricted to a minimum number of 32 proteins per leaf and have a maximum number of splits equaling three quarters the number of used features. We train N=600 ensembles on partitions of the data using 80% of class 1 (N1~40) and 10% of class 2 (N2~400). Thus, we obtain each protein's cross-validated score by the median score of these ensembles, excluding any runs where it was used in the training. When trained separately on experiment ($s_{exp}$) and sequence features ($s_{seq}$) and combined by the Euclidean distance, $s =$

$\sqrt[2]{\left(\max(s_{exp}) - s_{exp}\right)^2 + \left(\max(s_{seq}) - s_{seq}\right)^2}$, the final score $s$ shows a slightly better performance (AUC for recall LLPS 0.93 vs 0.91). Likely, this can be accounted to the relatively small training sets and that there are more sequence features than experimental features. The PSAP predictor could not be plotted, as its published output is does not contain cross-validated results of the training set.

#### Data processing

We create a gaussian null model for the measurement noise from replicates. To determine the shift from the diagonal of a protein when comparing two filtration conditions, we center the null model with a line fit to the upper edge of the data, assuming proteins do not elute less under higher pressure. To estimate the fraction of the proteome that is organized in liquid assemblies, we further constrain this by fitting the line through large complexes and proteins with transmembrane domains [23,24]. We sum up the shifts in the 30 nm differential pressure filtration and the 1.2x dilution experiments to quantify the 'liquid-like behavior'. This characterization returns similar results when including other conditions, however the centering of the null model is not as clear because fragile membrane bound organelles (e.g., Golgi apparatus and ER) start eluting more. We identify proteins in BMCs beyond a 2% false discovery rate (2.33 sigma of the null model), omitting any proteins with membrane annotation.

#### MS sample preparation and analysis

Low complexity samples: Preparation spin control, gravity flow and differential pressure filtration at 200 nm. Labeled using TMT-10plex (Thermo Fisher Scientific), analyzed by TMT-MS3 [25].

High complexity samples: Differential pressure filtration at 30 nm and 100 nm. Labeled using TMT-10plex, analyzed by TMTc+ [26]. Filtration of diluted cytoplasm. Labeled using TMTpro-16plex (Thermo Fisher Scientific), analyzed by TMTproC[27].

Sample preparation: Samples were prepared mostly as previously described [28]. Lysates were collected in 100 mM HEPES pH 7.2 and proteins were denatured by adding 2% SDS. To reduce disulfides, Dithiothreitol (DTT) (500 mM in water) was added to a final concentration of 5 mM (20 min, 60°C). Samples were cooled to RT, and cysteines were alkylated by the addition of N-ethyl maleimide (NEM, 1 M in acetonitrile) to a final concentration of 20 mM followed by incubation for 20 min at RT. 10 mM DTT (500 mM stock, water) was added at RT for 10 min to quench any remaining NEM. A methanol-chloroform precipitation was performed for protein clean-up, and the collected protein pellets were allowed to air dry. Samples were taken up in 6M guanidine chloride in 200 mM EPPS pH 8.5. Subsequently, the samples were diluted to 2 M guanidine chloride in 200 mM EPPS pH 8.5 for overnight digestion with 20 ng/μL Lys-C (Wako) at RT. The samples were further diluted to 0.5 mM guanidine chloride in 200 mM EPPS pH 8.5 and then digested with 20 ng/μL Lys-C and 10 ng/μL trypsin (Promega) at 37°C overnight.

The digested samples were dried using a vacuum evaporator at RT and taken up in 200 mM EPPS pH 8.0. The total material from each condition was labeled with tandem mass tags. TMT/TMTpro samples were labeled for 2 hours at RT. Labeled samples were quenched with 0.5% hydroxylamine solution. Samples from all conditions were combined into one tube, acidified to pH < 2 with phosphoric acid (HPLC grade, Sigma) and cleared by ultracentrifugation at 100,000 g at

4°C for 1 hour in polycarbonate tubes (Beckman Coulter, 343775) in a TLA-100 rotor. Supernatants were dried using a vacuum evaporator at RT. For a low complexity sample, dry samples were taken up in HPLC-grade water and stage-tipped for desalting [29] and resuspended in 1% formic acid (FA) to 1 µg/µL for mass spectrometry analysis. For high complexity samples, the supernatant was sonicated for 10 minutes and then fractionated by medium pH reverse-phase HPLC (Zorbax 300Extend C18, 4.6 x 250 mm column, Agilent) with 10 mM ammonium bicarbonate, pH 8.0, using 5% acetonitrile for 17 minutes followed by an acetonitrile gradient from 5% to 30%. Fractions were collected starting at minute 17 with a flow rate of 0.5 mL/min into a 96 well-plate every 38 seconds. These fractions were pooled into 24 fractions by alternating the wells in the plate[30]. Each fraction was dried and resuspended in 100 µL of HPLC water. Fractions were acidified to pH <2 with HPLC-grade trifluoroacetic acid, and stage-tipping was performed to desalt the samples. For LC-MS analysis, samples were resuspended to 1 µg/µL in 1% FA and HPLC-grade water, and ~1 µg of peptides were analyzed per 1 hour run time.

<u>MS analysis:</u> Approximately 1-3µg of the sample was analyzed by LC-MS. LC-MS experiments were performed with an nLC-1200 HPLC (Thermo Fisher Scientific) coupled to an Orbitrap Fusion Lumos (Thermo Fisher Scientific). For each run, peptides were separated on an Aurora Series emitter column (25 cm x 75 µm ID, 1.6 µm C18) (ionopticks, Australia), held at 60°C during separation by an in-house built column oven. Separation was achieved by applying a 12% to 35% acetonitrile gradient in 0.125% formic acid and 2% DMSO over 90 min for fractionated samples and 180 min for unfractionated samples at 350 nL/min at 60°C. Electrospray ionization was enabled by applying a voltage of 2.6 kV through a MicroTee at the inlet of the microcapillary column. As indicated in each proteomics experiment, we used the Orbitrap Fusion Lumos with a TMT-MS3 [25], TMTc+ [26], or TMTproC [27] as previously described.

Mass spectrometry data analysis was performed essentially as previously described [31]. The mass spectrometry data in the Thermo RAW format was analyzed using the Gygi Lab software platform (GFY Core Version 3.8) licensed through Harvard University. Peptides that matched multiple proteins were assigned to the proteins with the greatest number of unique peptides. TMT-MS3 [25], TMTc+ [26], or TMTproC [27] data were analyzed as previously described.

The mass spectrometry proteomics data have been deposited to the ProteomeXchange Consortium via the PRIDE [32] partner repository with the dataset identifier PXD029879. To access, use the username ‘’ and the password ‘TjFrM6Kh’.
